# Plastic responses to past environments shape adaptation to novel selection pressures

**DOI:** 10.1101/2024.05.06.592784

**Authors:** Sarah E. R. Coates, Aaron A. Comeault, Daniel P. Wood, Michael F. Fay, Simon Creer, Owen G. Osborne, Luke T Dunning, Alexander S. T. Papadopulos

**Author notes:** Corresponding Author: Alexander S.T. Papadopulos, **Email:**.

## Abstract

Phenotypic plasticity may pave the way for rapid adaptation to newly encountered environments. Although it is often contested, there is growing evidence that initial plastic responses of ancestral populations to new environmental cues may promote subsequent adaptation. However, we do not know whether plasticity to cues present in the ancestral habitat (past-cue plasticity) can facilitate adaptation to novel cues. Conceivably, this could occur if plastic responses are coincidentally optimal to both past and novel cues (i.e., are pre-adaptive) or if they are transferred to novel cues during adaptation. Past plastic phenotype values could also become fixed and genetically co-opted during adaptation to the new environment. To uncover the role of past-cue plasticity in adaptation, we tested gene expression plasticity responses of two parallel mine-waste adapted *Silene uniflora* populations and their closest coastal relatives. Plants were exposed to the past and novel-cues of salt and zinc, which revealed that during adaptation to mine-waste plasticity to salt diminishes. Despite this, our results show that ancestral plasticity to salt has a substantial impact on subsequent adaptation to zinc. For a third of genes that have evolved zinc plasticity in mine populations, salt plasticity has been transferred to the zinc response. Furthermore, a quarter of fixed expression differences between mine and coastal populations were similar to ancestral salt responses. Alongside evidence that ancestral plasticity to novel cues can facilitate adaptation, our results provide a clear indication that ancestral past-cue plasticity can also play a key role in rapid, parallel adaptation to novel habitats.

**Significance Statement:** The role of phenotypic plasticity in promoting adaptation is hotly debated, with conflicting evidence for the benefits of ancestral plasticity in newly encountered environments. Here, we present an alternative mode by which ancestral plasticity can promote adaptation. We investigated whether phenotypic plasticity towards environmental cues that are experienced only in ancestral habitats (past-cue plasticity) can significantly contribute towards rapid adaptation to completely distinct cues. We show that, in the maritime plant species, *Silene uniflora*, past-cue plasticity to salt has made a substantial contribution to rapid adaptation to heavy-metal pollution in newly encountered habitats. This phenomenon has broad implications for the capacity and predictability of species to persist in the face of anthropogenic environmental change.

## Introduction

Phenotypic plasticity is the ability of an individual genotype to produce different phenotypes in response to different environmental cues (1). Although plasticity can be adaptive (i.e., it increases fitness), the extent to which it can facilitate adaptation to novel habitats remains contested (2–8). One possible outcome is that plasticity moves a phenotype value closer to the optimum for the novel habitat (2, 4, 6). This initially plastic phenotype may then become genetically assimilated during adaptation (plasticity is canalised and no longer varies with environment), although the evidence is mixed for the likelihood of this process (3, 4, 8, 9). Additionally, during adaptation, selection may favour a change in the extent of plasticity in a phenotype (i.e., evolution of plasticity; 2, 10). Alternatively, initial plastic responses may be neutral (i.e., not under selection) or maladaptive (i.e., reduce fitness) and are reversed/reduced during adaptation (3, 9). Studies typically investigate these phenomena by focusing on whether the plastic change in an ancestral population in response to a novel environment (PC) moves trait values in the same direction as the evolutionary change (EC) in the derived population that follows adaptation (7–10).

Less attention has been devoted to the impact of ancestral plastic responses to past cues (i.e., those *only* experienced in the ancestral environment - here termed past-cue plasticity) on subsequent adaptation to different cues in the new environment. Past-cue plasticity may be lost during adaptation to a new environment due to being non-adaptive, or maladaptive. Alternatively, past-cue plasticity may potentiate novel adaptation by bringing trait values closer to the optimum for the new cue in the new environment. It is often thought that existing traits with one function may serve a new, beneficial purpose in new environments (11–14), i.e., the traits may be pre-adaptive. Existing phenomena suggest that tolerance to one stress may be pre-adaptive for additional stressors in plants - for example, co-tolerance has been observed between different heavy metals (15, 16) and salt and heavy metal-tolerance mechanisms may be shared (17–19). Despite the existence of co-tolerance of multiple stressors, there is little direct evidence to show that past-cue plasticity can be pre-adaptative for novel cues.

Here, we introduce a framework to establish the potential impact of past-cue plasticity on adaptation. We define three patterns which point to an influence of past-cue plasticity in directing post-adaptation trait values:

i. Pre-adaptive plasticity - Beneficial plasticity as a response to both past and novel environmental cues could pre-exist, for example due to pathways that facilitate co-tolerance to multiple stressors. This would be evident as similar phenotypic plasticity in the ancestral and descendent populations in response to both past cues and new cues, without any evidence of an evolutionary change (Figure 1A).
ii. Cue transfer – The plastic response to a past cue might be transferred to a new cue following adaptation. For example, genes expressed in response to light stress, but not heat stress, in the ancestral population may evolve a similar heat stress response in the descendent population. In practice, this can be characterised by determining if past-cue plasticity differs from PC *and* EC moves the descendent population’s plastic response to the new cue towards the past-cue plasticity value rather than to the PC value (Figure 1B).
iii. Co-option – Past-cue plasticity may bring a phenotype closer to the optimum value in the novel environment, resulting in the past-cue trait value becoming co-opted into constitutive genetic changes during adaptation (Co-option, Figure 1C). This is analogous to the genetic assimilation of plasticity to a single cue during adaptation (2, 4, 8). This can be identified if past-cue plasticity differs from PC and a constitutive EC during adaptation has taken the trait value closer to the ancestor’s past-cue value.

**Figure 1.**
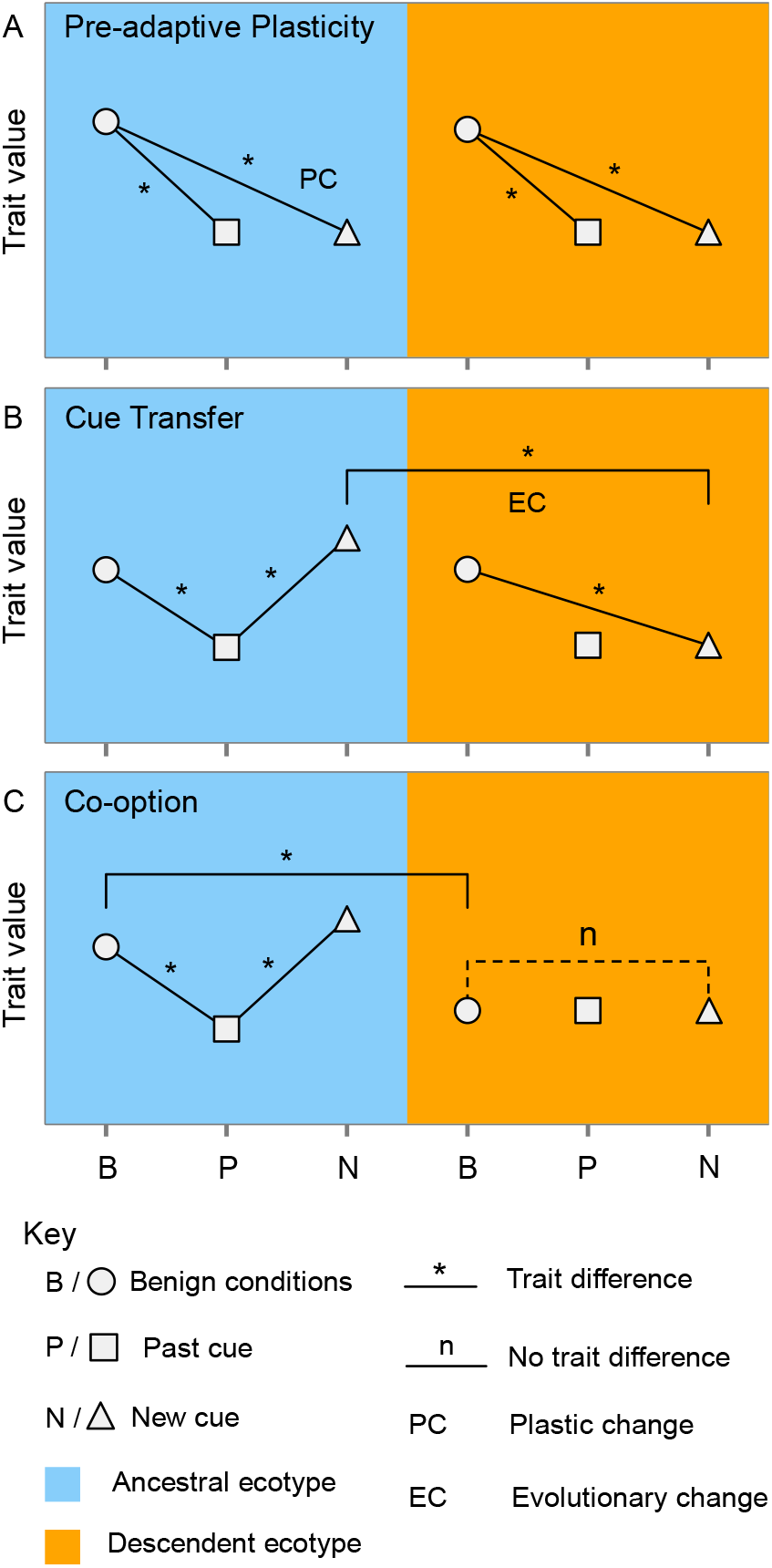
Framework of trait value comparisons for assessing whether plastic responses to past cues are co-opted or show cue transfer during adaptation to novel environments. Each panel shows the expected trait value changes in ancestral and descendent environments under different scenarios. Wherever no line is drawn, the trait can take any value. (A) Pre-adaptive plasticity, where ancestral and descendent populations share similar plasticity on exposure to past/new cues. (B) Cue transfer, where adaptation results in novel-cue plasticity resembling past-cue plasticity (C) Co-option, where adaptation to the novel environment results in constitutive expression shifting to become more similar to the ancestral population expression value in response to a past cue.

Using this framework, we assess the impact of past-cue gene expression plasticity during parallel adaptive evolution in *Silene uniflora*. In this generally coastal species, several populations have independently colonised and adapted to sites at abandoned industrial-era mines that are heavily contaminated with zinc (20). During adaptation, mine populations have evolved changes in gene expression that were facilitated by ancestral plasticity to the new environmental cue of high zinc concentrations. In this system, adaptation was characterised by evolutionary changes in the extent of plasticity and genetic assimilation (8, 20).

Coastal *Silene uniflora* are not exposed to high zinc levels, but they do grow in a challenging saline environment on cliff-tops and rocky shores. The degree of salt stress in this environment is spatially and temporally variable due to frequent changes in salt deposition rates from sea-spray and/or inundation (21, 22). Variability in environmental cues may enhance the evolution of plasticity (23, 24), therefore, we expect a high degree of gene expression plasticity in response to salt exposure in coastal populations. Using coastal populations as a proxy for the ancestors of mine populations, we tested whether gene expression plasticity to a past cue (salt) facilitates adaptation to a new cue (zinc) across two independent evolutionary replicates.

## Results and Discussion

To quantify the extent to which past-cue plasticity influences and is influenced by adaptation to new environments, we sequenced root transcriptomes of individuals from two pairs of coastal and mine-waste adapted *Silene uniflora* populations (Coast-W/Mine-W from Wales, 16.1km apart, and Coast-E/Mine-E from England, 25.6km apart) after hydroponic treatment with control and NaCl solutions (see Materials and Methods and SI Table S1). Additionally, we reanalysed transcriptomic data generated from a similar experiment which used the same populations, but grew plants in control and zinc solutions (8). This combination of experiments allowed us to determine; (i) the extent to which past-cue plasticity is lost during adaptation to a new cue, (ii) the role of pre-adaptive plasticity in adaptation, and (iii) the degree to which plastic responses can switch cues or be co-opted during adaptation.

### Adaptation to new cues alters the plasticity landscape

We quantified differential expression between coastal and mine populations in response to a past-cue (salt) and new-cue (zinc) to compare the ancestral and descendent responses to both cues. The coastal populations shared 957 salt-plastic genes with the same direction of expression change (more than expected by chance: randomisation test, p = <0.00001, SI Table S2; Figure 2A), which is roughly half the total number of salt plastic genes in each individual population (Coast-W = 2,078, Coast-E = 1,676; Figure 2A). Only 155 genes were salt-responsive in both mine populations (randomisation test, p = <0.00001, SI table S2; figure 2A). This demonstrates a substantial and parallel loss of plasticity (86.21%) in response to salt stress following adaptation to the mine environment. Although plasticity is reduced, the pattern of expression in response to salt within mines resembles that of their coastal ancestors; 85% (132) of the 155 shared mine salt-plastic genes were also plastic in response to salt in both coasts. Upon exposure to salt stress, the proportion of the whole transcriptome that was differentially expressed in both coast/mine populations was quite modest (4.14% for coasts and 0.67% for mines) (Figure 2B) when compared to the transcriptome-wide zinc-stress response of coastal populations (47.34%) (8). We compared the functions of enriched Gene Ontology (GO) terms between coastal and mine salt-plastic genes. The breadth of salt-plastic gene functions reduced during adaptation to zinc, although some functions remain shared between coasts and mines (Figure 2C, SI Tables S3-S5, SI datasets S1, S2). Environments with consistent, rather than variable, cues are expected to select for reduced plasticity (24). Therefore, exposure to consistently low salt concentrations in the mine environment may have selected for reduced salt plasticity, but some ability to tolerate variable salinity has been retained despite adaptation to zinc. Such retention of plastic responses to past cues from the ancestral environment may underpin the dominance of plastic changes over genetic adaptations when ancestral environments are recolonised, as found by Ho *et al*. (25).

**Figure 2.**
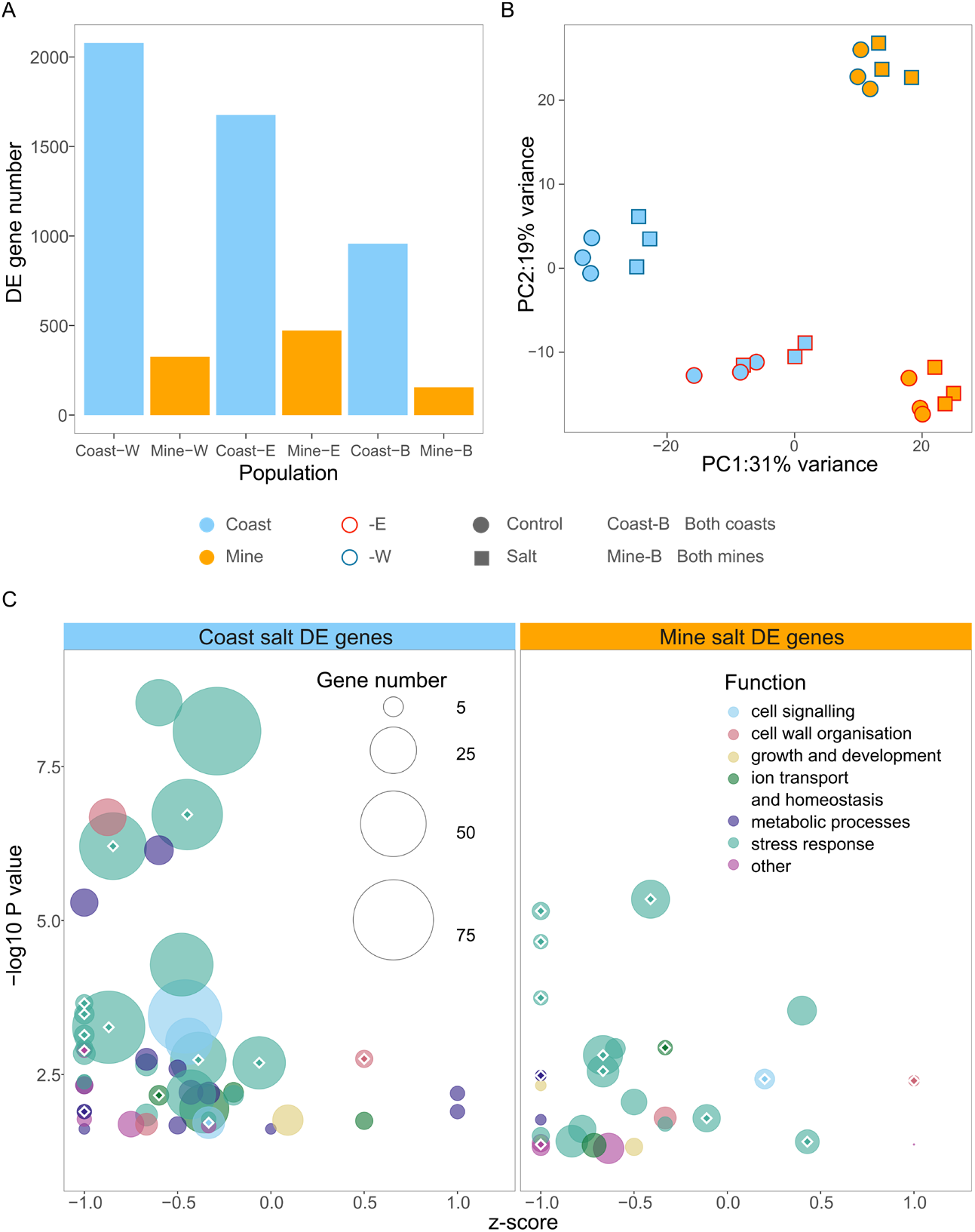
The impact of new-cue (zinc) adaptation on past-cue (salt) response plasticity. (A) Total number of salt-induced differentially expressed (DE) genes in each population and those genes with the same direction of expression change in both coasts/both mines (Coast-B/Mine-B). (B) PCA of variance-stabilised transformed counts of 30,714 genes in each population in control and salt treatments (C) Bubble plot showing the z-score of coastal and mine genes differentially expressed in salt within significantly enriched GO categories (*n* upregulated – *n* downregulated/total *n* in each GO category) against the negative log transformed p-value for each GO term. 1.0 = 100% upregulated genes, -1.0 = 100% downregulated genes, 0.0 = 50% up/downregulated genes. Bubbles scale with the number of genes in each GO category and colours represent sets of broader common functions. The 14 common significant GO terms are shown with white outlined diamonds.

In line with Wood *et al*. (8), the reanalysed zinc experiment included 10,933 zinc-plastic genes shared by both coastal populations. Mine-adapted populations shared 143 zinc-plastic genes (SI Table S2) with 91 undergoing an evolutionary change in plasticity to zinc (63%). In control treatments, 124 genes were differentially expressed between both pairs of coastal and mine populations, showing a pattern of constitutive evolutionary change during adaptation. These two sets of genes (143 with zinc-plasticity and 124 with constitutive evolutionary change) are likely to be involved in adaptation to the mine environment across the independent replicates (8).

### No evidence for pre-adaptive plasticity

To test for evidence of the role of pre-adaptive plasticity during adaptation to novel stressors we quantified the number of genes which had significant expression changes that were similar across both treatments, and both mine and coast population pairs. We found that there were no genes with this pattern, demonstrating that pre-adaptive plasticity has not played a role during adaptation to this novel environment. Plastic responses to one stressor which are coincidentally also beneficial to another stressor might be expected, as co-tolerance has evolved between pressures that have similar impacts on plant physiology (15–19) or for chemically similar ions, such nickel and lead (15) or zinc and nickel/cobalt (16). Both salt and heavy metals produce reactive oxygen species and some molecular mechanisms are likely to alleviate the impacts of both stressors (e.g., antioxidants; 17, 18). However, it is clear from this result that salt tolerance does not automatically and instantaneously confer zinc tolerance. Some species are known to possess both salt and heavy metal tolerance in the same populations when they occur in habitats with both stressors present (19, 26, 27). Our result suggests that adaptation to both stressors is required in these cases rather than adaptation to one stressor being pre-adaptive for the other. This indicates that past-cue plasticity is not strictly pre-adaptive, but this may depend on the similarity (e.g., chemically) between the past and new cues encountered by the species.

### Past-cue plasticity is transferred to novel cues during adaptation

We tested for signals of cue-transfer during adaptation by quantifying the number of coastal salt-plastic genes that underwent a change in zinc plasticity during adaptation and for which mine zinc plasticity matched the direction of the coastal salt response. Cue-transfer occurred for almost one third of the genes with evolved zinc plasticity (30.77%, 28/91, Figure 3A), demonstrating that the repurposing of past-cue plasticity during adaptation to a new cue can play a substantial role in adaptation (SI Figure S1). Many molecular pathways are commonly involved in alleviating the consequences of different environmental stresses in different species. In this case, a large component of adaptation may simply be modifying the sensitivity of the pathways to the new stressor. Indeed, several of our putative adaptive genes have been implicated in tolerance to both salt and heavy metals, including those involved in signalling pathways (see section below).

**Figure 3.**
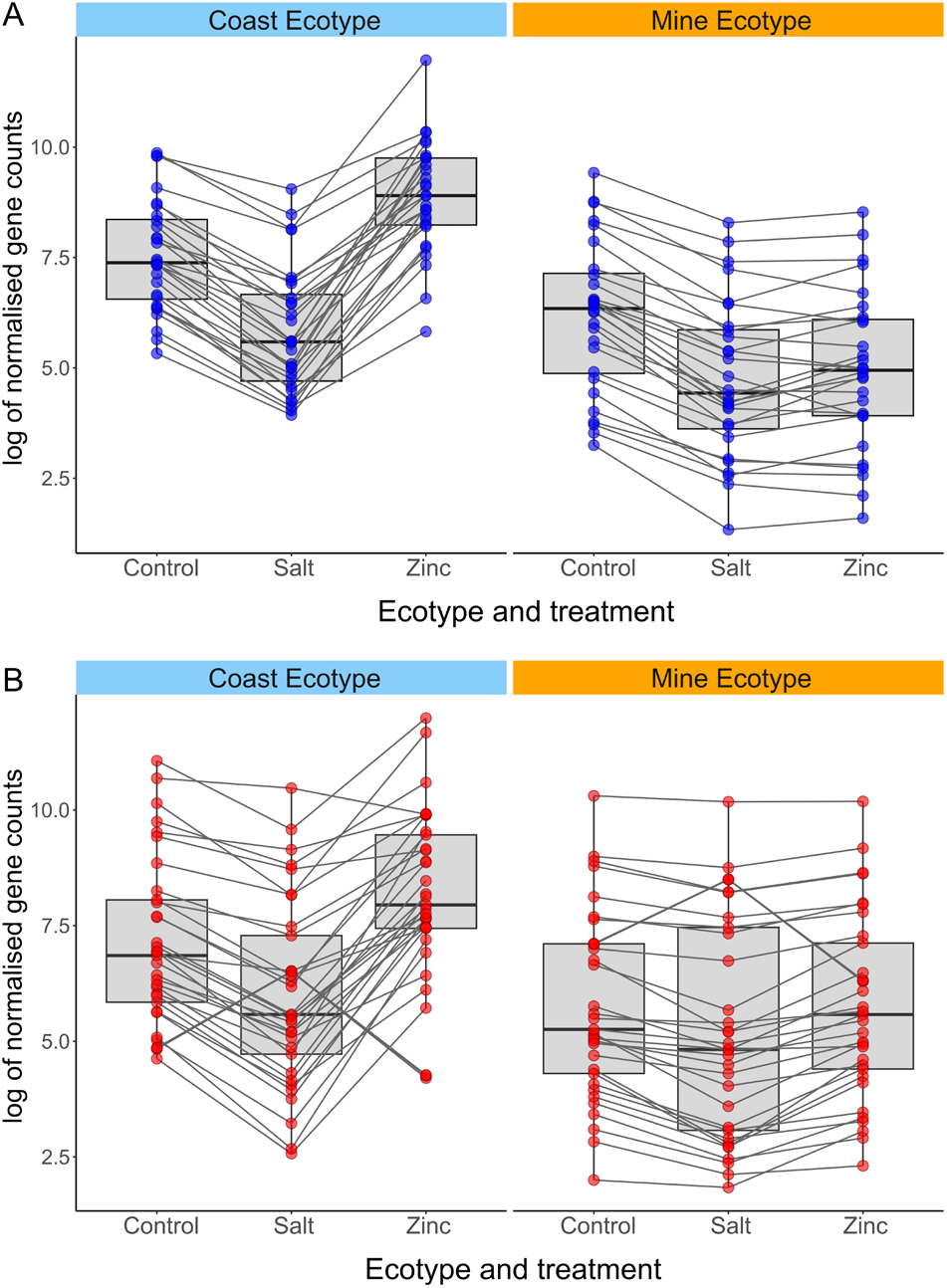
Box and line plots showing the natural log of normalised mean gene expression counts in coast and mine ecotypes (both Welsh and English populations) across control, salt, and zinc treatments, for different gene sets. (A) Cue transfer genes (*n* = 28) and (B) Co-opted genes (*n* = 33). Points represent individual genes and lines show how mean expression counts differ between each treatment for each gene.

Many studies have focused only on changes in plasticity in response to the same cue in ancestral and adapted populations (3–5, 7–9). Under the frameworks used in these studies, genes that have undergone cue transfer would be classified as undergoing reversion after adaptation. Thus, plasticity in these genes would appear to be maladaptive/non-adaptive in the ancestor, when in fact they harbour adaptive plasticity for novel cues within the past-cue response. Consequently, the role of ancestral plasticity in facilitating adaptation to novel environments may be greater than previously estimated.

### Past-cue plasticity is co-opted during adaptation

To provide evidence for co-option we determined the number of coastal salt-plastic genes with no significant zinc response plasticity in mine plants which did display a constitutive evolutionary change matching the direction of the salt response. In other words, genes for which ancestral plasticity has been lost, but the zinc-adapted trait value is close to the ancestral salt-response. In total, 26.61% (33/124) of genes with a constitutive evolutionary change (genes differentially expressed between mine and coastal plants in the control conditions) had expression patterns consistent with co-option (Figure 3B, SI Figure S2). This is a similar proportion to those undergoing cue transfer, suggesting that co-option is almost as likely as cue-transfer during novel adaptation. Previously, Wood *et al*., (8) found that close to 50% of the genes with constitutive evolutionary changes had undergone genetic assimilation of ancestral zinc plasticity. The co-opted genes detected here are mutually exclusive of the genetically assimilated genes, but are from the same larger set (i.e., they all have constitutive evolutionary changes in expression). Taken together, close to three quarters of these constitutive expression changes observed in *S. uniflora* appear to have been facilitated by ancestral plasticity of some kind. This suggests that ancestral plasticity that brings expression values closer to the optimum for the new environment are likely to be channelled into fixed expression responses during novel adaptation, regardless of whether this plasticity was in response to past or novel cues. Our results show that to understand the role of plasticity more fully during adaptation, it is paramount to test responses to environmental cues found in both ancestral and novel environments.

### Evidence for co-functionality in cue transfer and co-opted genes

Although many of the precise functions of genes undergoing cue transfer and co-option are unknown, several have been implicated in tolerances to both salt and zinc in different species (SI Dataset S3, S4) suggesting they have switched or gained functions during adaptation. Genes encoding chalcone synthase-like proteins (CHS2) were detected among both cue-transfer and co-opted genes - chalcone synthase is a key structural enzyme in the flavonoid biosynthetic pathway (28, 29) and has a potential role in the chelation of heavy metals such as copper, lead, cadmium and nickel (30, 31). Chalcone synthases have also been linked to salinity tolerance (32). Genes encoding enzymes with transferase activities were also found in both sets (Cue-transfer - a glutathione-S-transferase; Co-option - a UDP-dependent glycosyltransferase). These transferases belong to gene families which are often implicated in both salt and heavy metal stress responses (33–36). Strikingly, one co-opted gene *Two Pore Channel 1* (*TPC1*), is known to be involved in rapid systemic signalling under salt stress in *Arabidopsis thaliana* (37), is a candidate for zinc tolerance in *Thlapsi caerulescens* (38) and has been strongly implicated in playing a key role during parallel adaptation to serpentine soils in *Arabidopsis arenosa* (39).

Cue transfer and co-opted genes were also enriched for GO terms linked to osmotic, oxidative and other abiotic stresses linked to heavy metal responses (SI Table S7). These results support the hypothesis that co-functionality is present between zinc and salt which may be due to both stressors having overlapping impacts on physiology (17). It may be the case that the more similar the past and novel cue, the more likely it is that past-cue plasticity will influence adaptation. This co-functionality might explain why we see relatively little indication that there are strong trade-offs between zinc and salt tolerance under these controlled, stable conditions. Most genes with cue transfer retained significant salt plasticity in mine populations (Figure 3A) and, unlike the widespread maladaptive transcriptomic response to zinc in coastal plants, expression profiles of mine plants did not shift in response to salt any more than the coastal plants. The observed responses under the stable and competition-free conditions of our experiment suggest that many mine adaptations might be conditionally neutral (or even beneficial) in the salt treatment, but these changes might be more disadvantageous in variable, high-competition conditions of wild coastal habitats.

## Conclusion

The role of plasticity in adaptation has become increasingly disputed with evidence both for and against reinforcement of plasticity and genetic assimilation in the process. We leveraged instances of parallel adaptation to a recently created novel environment to test the contribution of plasticity to cues from both the new and ancestral environment. Overall, three quaters of the fixed expression differences between ancestral and derived populations can be linked to ancestral plasticity to the past or new cue. Our experiments demonstrate that there is a substantial contribution of ancestral plasticity to both the evolution of new plasticity in expression and canalised expression levels during adaptation.

## Materials and Methods

### Plant sampling and hydroponic experiment

We studied four populations; Coast-W, Mine-W, Coast-E and Mine-E corresponding to WWA-C, WWA-M, ENG-C, ENG-M in Papadopulos *et al*. (20) and S1, T1, S2, T2 in Wood *et al*. (8). An experiment to assess zinc associated gene expression change was carried out as described in Wood *et al*. (8). We then carried out a near-identical experiment to determine salt (NaCl) associated expression change. Three individuals per population were cloned via mist-propagation and acclimated to deep water hydroponic tanks containing Hoagland’s nutrient solution. After one week, solutions were replaced with either the same solution as a control or the solution plus 0.1M NaCl (three clones per individual per treatment, see SI methods). After eight days, root tissue from the clones of each individual was pooled and total RNA was extracted with a Qiagen RNeasy plant kit (See SI Methods for more detail). The 24 pools were sequenced on an Illumina Novaseq platform by Macrogen Genomics Europe. The read length was 100bp (mean insert size = 101 bp) and the total number of reads per sample was between 40.2 and 43.8M (SI Table S8).

### Transcriptome assembly and expression counts

Raw reads from both the salt and zinc datasets (8) were quality checked and trimmed to remove adapters (see details in SI Methods and Table S8). We used STAR version 2.7.10a (40) to map the trimmed reads to the *S. uniflora* reference genome (see SI Methods). The transcriptome was then assembled against the reference genome annotation (41) using StringTie v2.2.0 (42). We generated separate gene expression count matrices for the salt and zinc experiments (41,603 genes in each) using the StringTie prepDE.py3 script (SI datasets S5 and S6).

### Differential expression analysis

We used the R package DEseq2 v1.40.0 (43) to analyse gene expression data. We filtered the zinc and salt datasets to remove sample counts of <10 and combined them to generate cross-experiment data. To ensure cross-experimental comparability, we filtered all results by genes with no significant differential expression in control conditions between the two experiments, leaving 23,093 genes for further analysis (see SI Figure S4 and SI Methods for more detail). We conducted principal components analyses with the R package prcomp for the salt (30,714 genes, Figure 2B) and combined experiment datasets (30,178 genes, SI Figure S3) using variance stabilised transformed counts.

We used two models in DEseq2 to test for differential expression with input gene counts and phenotype data (SI datasets S5, S6 and S7). The first consisted of a single combined factor of *Population + Treatment* to compare within-treatment gene expression between populations and within-population expression between experiments. The second compared within-population gene expression between salt and control or zinc and control treatments, with the formula: *∼ Population + Population:Individual + Population:Treatment* (details in SI methods).

### Differential expression contrasts for hypothesis testing

We used multiple combinations of differential expression contrasts to determine the impact of novel adaptation on past-cue plasticity and to provide evidence for processes of co-option, cue transfer and pre-adaptive plasticity (Figure 1). Coast or mine salt/zinc plastic genes were those that were differentially expressed between control and salt/zinc treatments in the same direction in both coast or both mine populations. Genes with evolved plasticity to zinc were defined as those differentially expressed between both mine and coastal populations in the zinc treatment *and* had zinc plasticity in mine populations. Genes with evolved constitutive expression change were defined as those that were differentially expressed between each coast and mine in control conditions as in Wood *et al*. (8).

To test for pre-adaptive plasticity, we searched for genes with the same plasticity to salt/zinc, *and* shared salt and zinc plasticity in coastal plants (SI Figure S5A). To test for cue transfer, we identified the coastal salt plastic genes that were differentially expressed between salt and zinc treatments in coastal plants *and* had evolved plasticity to zinc in the mine populations (SI Figure S5B). To test for co-option, we searched for genes involved in coastal responses to salt that also had constitutive evolved expression changes, *were not* differentially expressed between control and zinc treatment in mines and *were* differentially expressed between salt and zinc treatments in both coastal populations (SI Figure S5C). See SI methods for more details.

### Functional analyses

The function of genes within the sets of interest was determined using the *Silene uniflora* reference annotation (41). We also conducted Gene Ontology (GO) enrichment using topGO v2.52.0 (see SI methods for more details).

## Acknowledgments

This work was funded by a Natural Environment Research Council standard grant NE/R001081/1 (awarded to ASTP) and the NERC Envision Doctoral Training Program NE/L002604/1 (awarded to SERC). We thank N. Welsby and H. Simpson for technical/greenhouse support. LTD was supported by a NERC Independent Research Fellowship (NE/T011025/1). Raw sequencing data is available via the Short Read Archive (PRNJXXXXXXX).

## Author Contributions

ASTP conceived the research and supervised the project along with AAC, SC and MFF. ASTP and DPW designed the experiment, with input from SERC. SERC conducted experimental and laboratory work. SERC analysed the data with input from ASTP, AAC, DPW, SC, MFF, LTD and OGO. SERC and ASTP wrote the manuscript and all authors provided comments on the manuscript before submission.

