## Supplemental information for "Plastic responses to past environments shape adaptation to novel selection pressures"

**Supporting Information Text**

**Supplementary Methods**

**Experimental design and hydroponics**

To analyse salt associated gene expression changes, we used three individuals from each of four populations. The individuals were either from existing *S. uniflora* living collections (n = 4, germinated as in Wood *et al.* (1)) or more recently germinated individuals (n = 8). All samples were grown from seed collected as part of a previous study (2) - population codes in this study: Coast-W, Mine-W, Coast-E, Mine-E; correspond to WWA-C, WWA-M, ENG-C and ENG-M in Papadopulos *et al.* (2); and S1, T1, S2, T2 in Wood *et al.* (1), respectively. One Mine-W individual and one Mine-E individual were the same plant as sampled in Wood *et al.* (1). Newly germinated plants were transferred to 1.5L pots three months after germination and grown on under ambient conditions for c. eight months. Cuttings were taken from three individuals per population and were rooted using mist propagation for two weeks with 12L di$H_2O$ and a further two weeks with 12L di$H_2O$ containing 1.92g Hoagland's nutrient powder. After four weeks of propagation, an equal proportion of cuttings from each population (n = 6) were transferred to six deep water hydroponic growth tanks filled with 0.16%w/v Hoagland's nutrient solution and allowed to acclimate for one week. The Hoagland's nutrient solution was made up by dissolving 1.28g of Hoagland's number two powder in 0.8ml di$H_2O$ per tank and adding KOH to adjust the pH to 5.5. The solution was diluted in each tank by adding 7.2L di$H_2O$ per tank, taking the total volume per tank to 8L. After acclimation, three tanks received fresh Hoagland's solution alone and three tanks received 0.1M NaCl dissolved in 0.16%w/v Hoagland's nutrient solution (three clones per individual in each treatment). Similar concentrations of NaCl elicit changes in phenotype in plant species such as other *Silene* (3, 4) and the halophyte *Armeria maritima* (5, 6). After eight days of these treatments, root tissue from the clones of each individual were pooled within treatments, flash frozen in liquid nitrogen and stored at -80°C. Pools of root tissue were homogenised and total RNA was extracted from 70-100mg tissue using a Qiagen RNeasy plant kit with an elution volume of 50μl. Tissue disruption with stainless steel beads and sand was carried out via the Tissuelyser Lt machine with a cooled head at 50Hz for two minutes. RNA sequencing (100bp, paired end) of the resulting 24 root samples (three individuals per population) was carried out using TruSeq stranded mRNA (Illumina) library preps and run on an Illumina Novaseq by Macrogen Genomics Europe. The total number of reads (paired, forward and reverse) was between 40.2 and 43.8 million per sample (SI table S7).

**Read processing and mapping**

Quality control information was generated with FastQC (7) and summarised using multiQC software (8). Reads were trimmed and filtered using Trimmomatic, with the options of: -phred33 ILLUMINACLIP:${MYADAPS}:2:30:10 LEADING:3 TRAILING:3 SLIDINGWINDOW:4:10 MINLEN:50 (9). MYADAPS='/apps/genomics/trimmomatic/0.39/adapters/TruSeq3-PE-2.fa'. The reference genome *S. uniflora* (10) was indexed, and reads of each individual sample mapped to the reference genome using STAR v2.7.10a (11). To complete the mapping step, STAR v2.7.10a (11) was run with the following options: "outSAMtype BAM SortedByCoordinate; outSAMstrandField intronMotif; outSAMattributes NH HI AS nM XS". Reference-based transcriptome assembly was carried out in StringTie v2.2.0 (12) using the reference annotations for *S. uniflora* and the following options: -e -B -p 10. We used the transcriptome assembly to generate a matrix of gene counts for 41,603 genes for both salt and zinc experiments in the format for downstream analysis by running the StringTie python script, prepDE.py3, with the -g option (SI datasets S5 and 6).

**Differential expression analyses**

We used the R package DEseq2 v1.40.0 (13) to analyse our gene expression count data and test for differential expression. We filtered the zinc and salt count datasets to remove sample counts of <10 and combined them to generate cross-experiment expression data. We conducted principal components analyses with the R prcomp function for the salt experiment alone (30,714 genes, figure 2B) and for the experiments combined (30,178 genes, SI figure S3) using variance stabilised transformed counts. The control treatments for both salt and zinc experiments were tightly clustered in the PCA (SI figure S3). This suggests the two separate experiments were highly consistent and, thus, could be combined. To ensure comparability between the experiments for expression comparisons, we created a set of genes that were not differentially expressed between the control conditions for the two experiments (SI figure S4). Only genes for each population that were not differentially expressed between controls were included. After this filtering, 23,093 genes were available for further analysis (~56% of the total number assembled).

We analysed the three filtered expression matrices and corresponding phenotype data (SI dataset S7), using DeSeq2's built-in models to identify genes that were differentially expressed. We applied two differential expression models to quantify differential expression in three ways: (i) between population differential expression within treatments (e.g. Coast-W control vs Mine-W control), (ii) between experiment and treatment conditions within populations (e.g. Coast-W salt experiment control vs Coast-W zinc experiment control and salt vs zinc comparisons); and (iii) between treatments, within populations and experiments (e.g. control vs salt in Coast-W). The first model to test (i) and (ii) consisted of a single combined factor of *Population+Treatment*. The second model examined (iii) using the formula: ~ *Population* + *Population:Individual* + *Population:Treatment*.

**Framework of differential expression contrasts for hypothesis testing**

We used multiple combinations of differential expression comparisons to determine the impact of novel adaptation on past-cue plasticity and to provide evidence for processes of co-option, cue transfer and pre-adaptive plasticity (Figure 1). We first categorised genes into distinct groups. Coast or mine salt/zinc plastic genes were those that were differentially expressed between control and salt/zinc treatment in the same direction in both coast or in both mine populations. The genes that have newly evolved plasticity to zinc were those that were differentially expressed between both mine and coastal populations in the zinc treatment *and* between control and zinc in mine populations. Genes that have evolved constitutive expression changes were defined as those that were differentially expressed in the same way between each coast and mine in the control conditions as in (1).

To test for pre-adaptive plasticity, we searched for genes with shared mine salt and zinc plasticity (i.e., mine responses to both treatments were the same), *and* shared salt and zinc plasticity in coastal plants (SI figure S5A). To test for cue transfer, we identified the coastal salt plastic genes that were differentially expressed between treatments in coastal plants (i.e., ancestral salt and zinc responses were not the same) *and* which had also evolved shared plasticity to zinc in the mine populations (SI figure S5B). To test for co-option, we searched for genes involved in coastal responses to salt that also had evolved constitutive expression changes, were *not* differentially expressed between control and zinc treatment in mines and *were* differentially expressed between salt and zinc treatments in both coastal populations (supplementary figure S5C).

The significance of the number of differentially expressed genes in response to salt and zinc shared across population pairs was analysed using a randomisation test. The test consisted of 10,000 replications of random draws of differentially expressed genes from the total number of filtered genes (23,093). An empirical p value was calculated for each randomisation test, which was calculated as the frequency of randomisations producing overlaps that were more than or equal to the observed number divided by the number of replications. We also determined the maximum overlap for each randomisation for comparison with the observed overlap.

**Functional analyses**

We determined the function of genes within the comparison sets of interest using *Silene uniflora* reference annotations (10). We conducted GO ontology enrichment to detect functional enrichment within our gene sets of interest using topGO v2.52.0 (14). We conducted this analysis on genes plastic to salt in both coasts and mines (tables S2-S4), and for cue transfer and co-opted gene sets (tables S5 and S6). To determine how the enriched GO terms differed between mines and coasts in salt plastic genes, we determined the *z*-score (z = sum(+ve LFC genes) – sum(-ve LFC genes)/ total n of genes in GO category) for each gene within a significant enriched GO (x axis, Figure 2C). We based this calculation from the *z*-score calculation described in the R

GOplot package (15). We highlighted GOs by broader shared functional categories that were defined based on the descriptions of the GO terms (Figure 2C, tables S2-4). This gives an indication of the patterns of expression change within the enriched GO term. For the sets of genes with cue-transfer and co-option patterns, we searched for enriched GO terms linked to salt/osmotic stress and heavy metal stress to find co-functionality between these stress responses. We also studied the functional annotations themselves to look for genes with potential salt and heavy metal stress co-functionality.

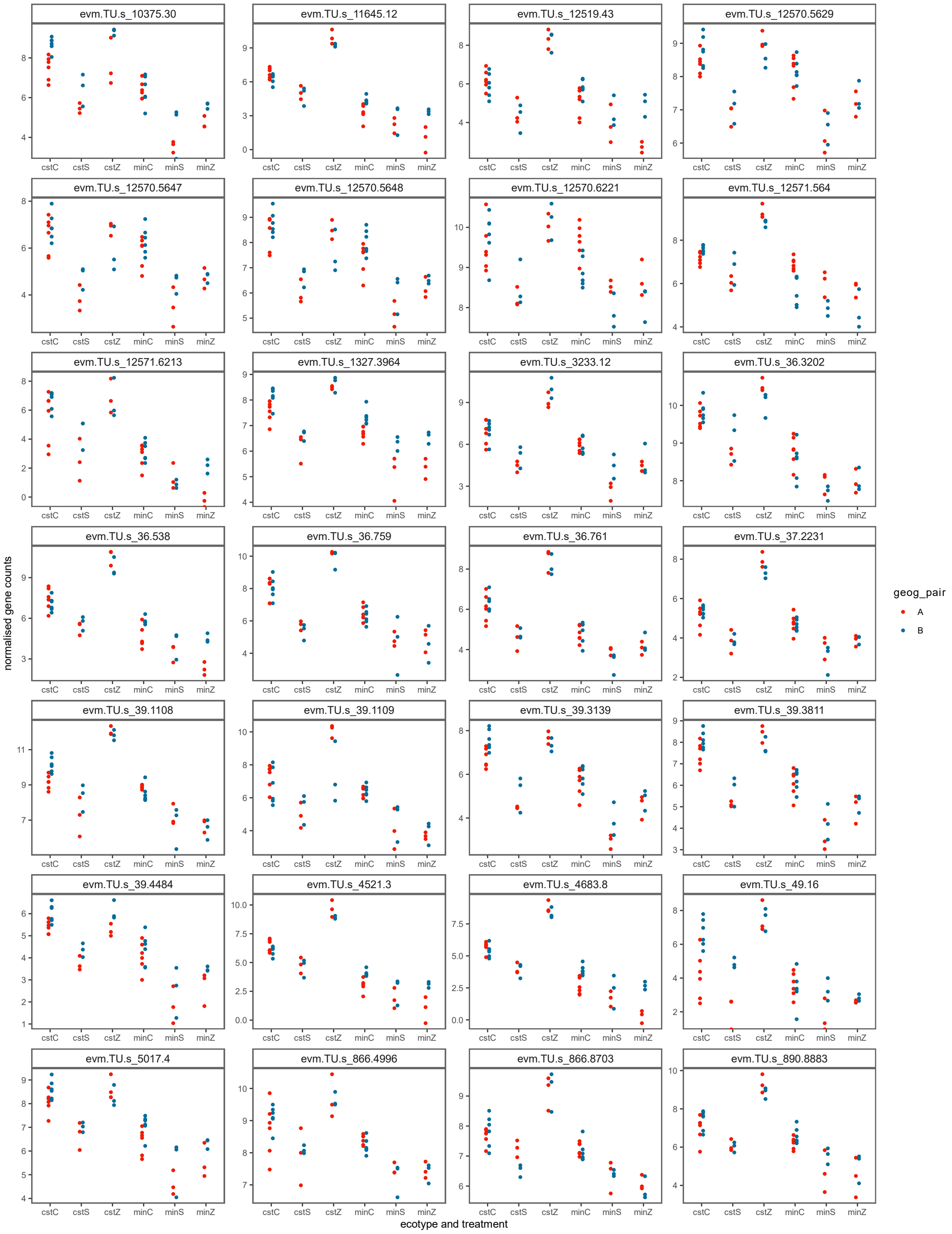


**Fig. S1. Gene by gene expression patterns for 28 Cue transfer genes.** Normalized expression counts across all 6 ecotype-treatment combinations for each gene with evidence of co-option. Each point represents a sequenced individual and the different colours represent each population studied. The control treatments contain 6 individuals per population as zinc/salt experimental samples have been combined and the zinc and salt treatments have 3 individuals per population.

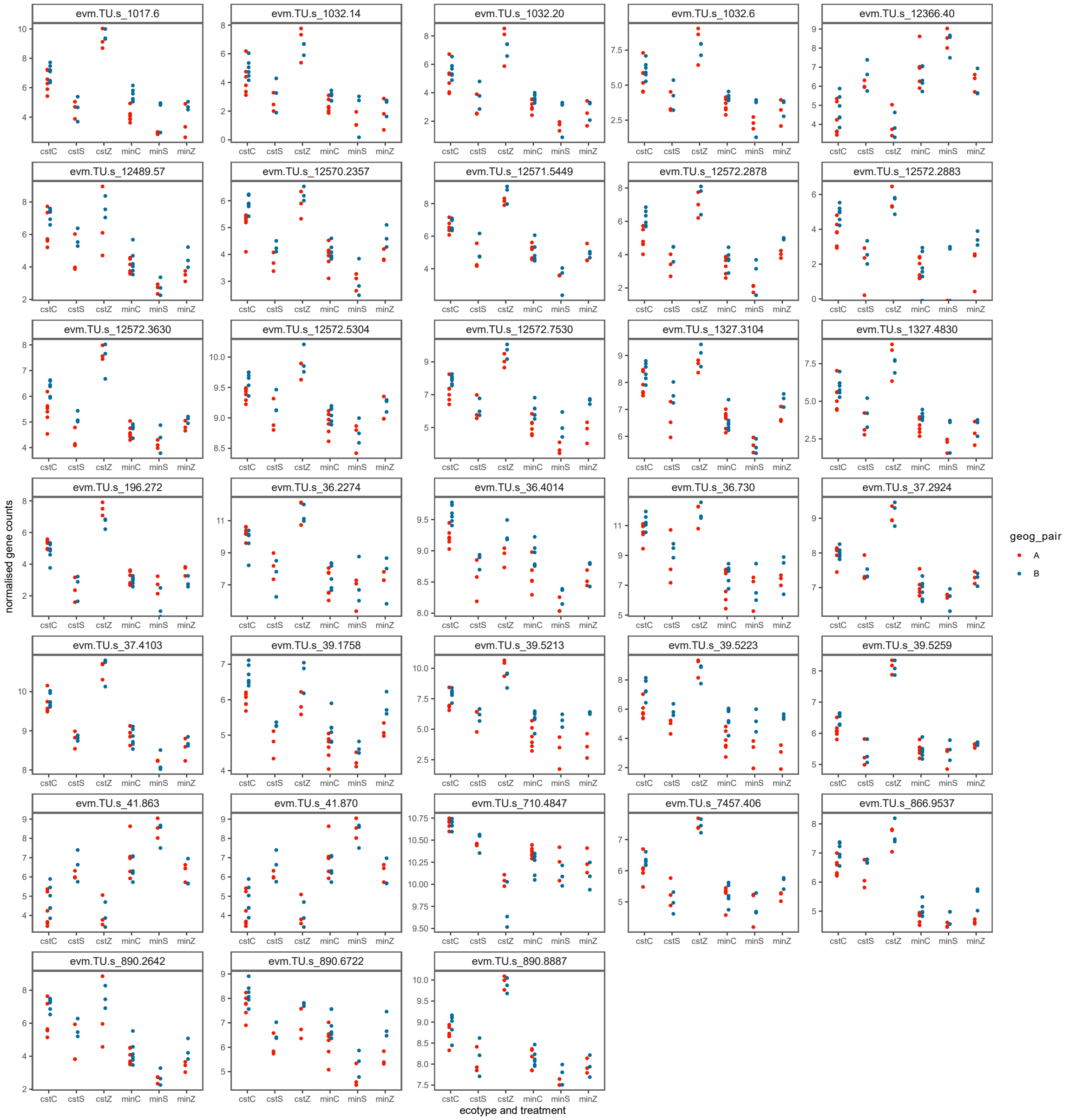


**Fig. S2. Gene by gene expression patterns for 33 Coopted genes.** Normalized expression counts across all 6 ecotype-treatment combinations for each gene with evidence of co-option. Each point represents a sequenced individual and the different colours represent each population studied. The control treatments contain 6 individuals per population as zinc/salt experimental samples have been combined and the zinc and salt treatments have 3 individuals per population.

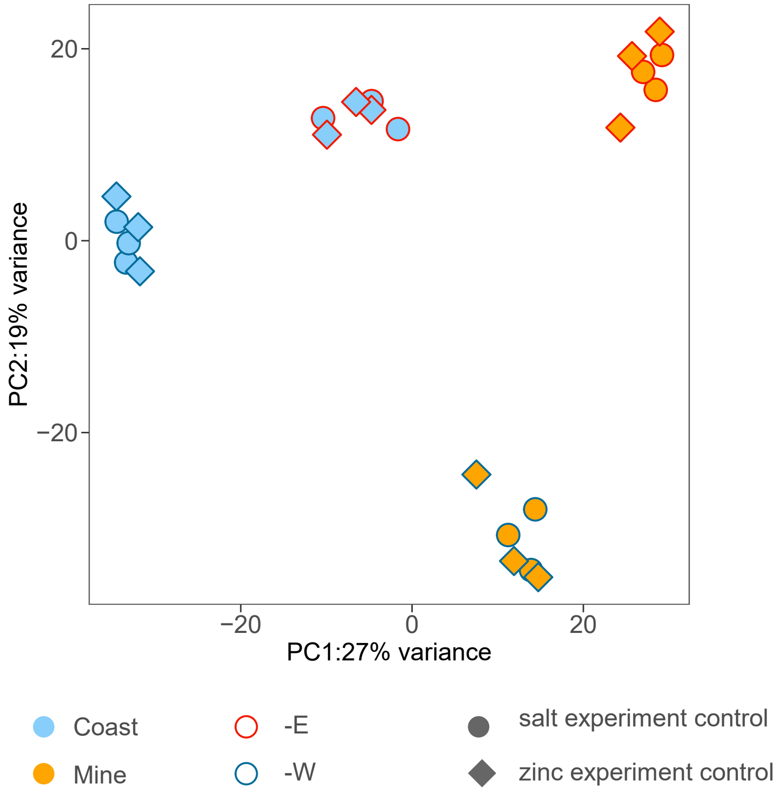


**Fig. S3.** Principle components 1 and 2 from PCA of control treatments from salt and zinc experiments.

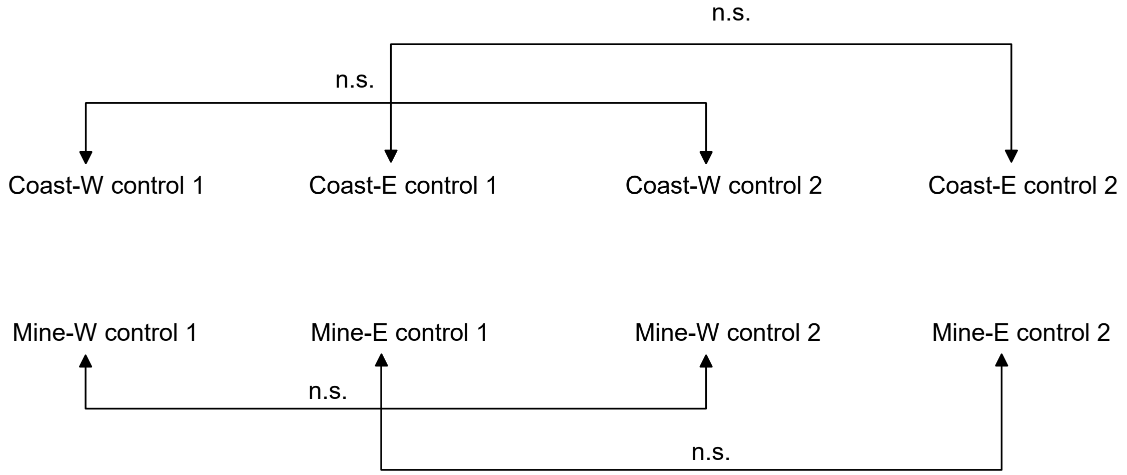


**Fig. S4** Set of differential expression contrasts used for ensuring cross-experimental comparability (n = 23,093). n.s.= no significant differential expression, Salt experiment = 1, Zinc experiment = 2.

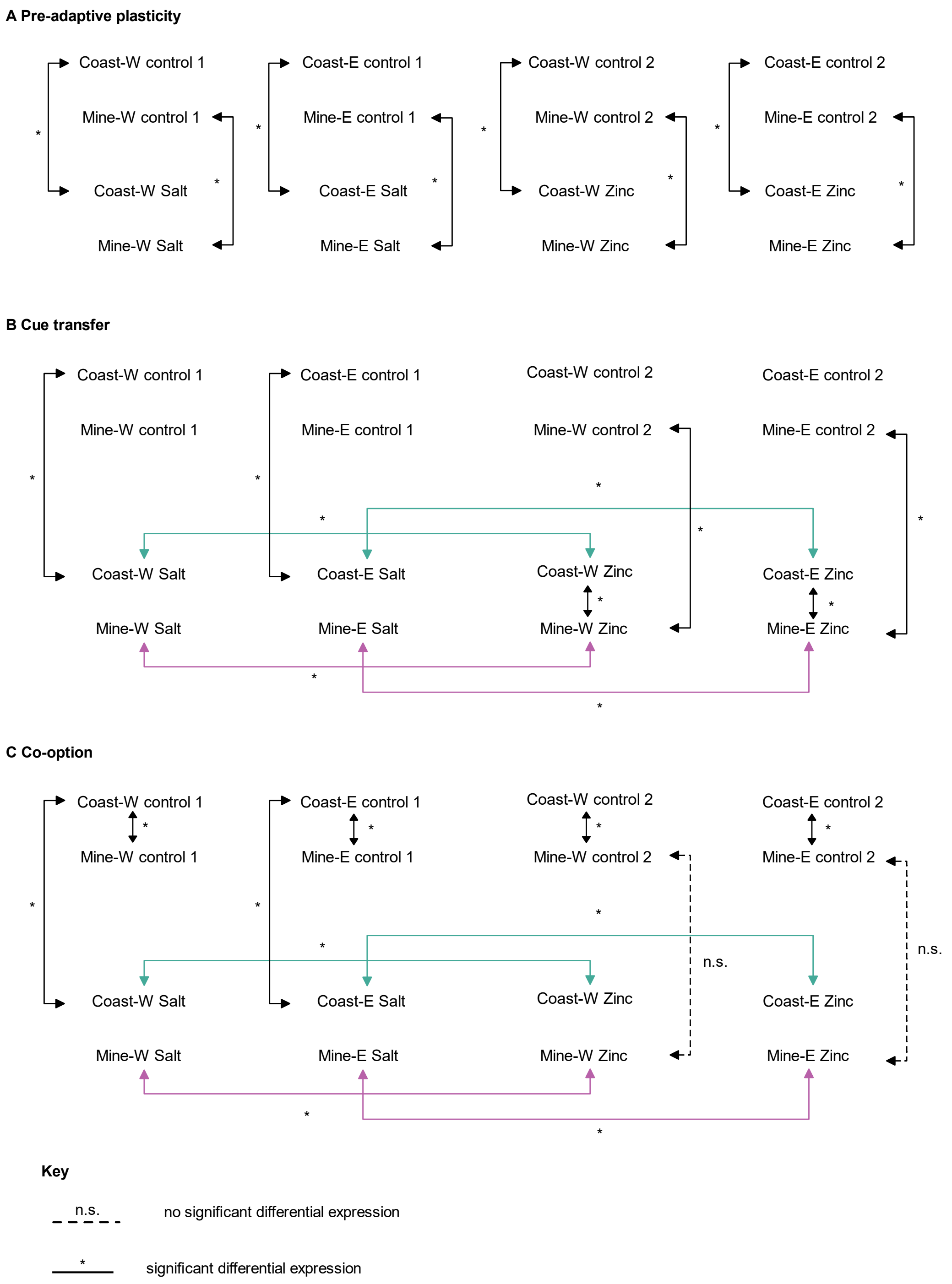


**Fig. S5.** Differential expression comparisons network for (A) pre-adaptive plasticity (n=0), (B) cue transfer (n=28) and (C) cooption (n = 33). Shared line colours to mark the contrasts denote the sets that were filtered to those only with the same direction of expression change across the set.

**Table S1.** Population coordinates and locations for sampled *Silene uniflora* populations. Included are the population names used in both Wood *et al.* (1) and Papadopulos *et al.* (2) The two letter population codes used in sequencing are also listed.

| Population name | Name in Papadopulos *et al.* 2021 | Name in wood *et al.* 2023 | Latitude | Longitude | Country | Location | Population code for sequencing |
|---|---|---|---|---|---|---|---|
| Coast-W | WWA-C | S1 | 52.394825 | -4.0939136 | Wales | Aberystwyth | SA |
| Mine-W | WWA-M | T1 | 52.331608 | -3.887207 | Wales | Grogwynion | GR |
| Coast-E | ENG-C | S2 | 51.323284 | -3.0169961 | England | Brean Down | BD |
| Mine-E | ENG-M | T2 | 51.256935 | -2.6509495 | England | Priddy Pools | PP |

**Table S2.** Sets of shared salt/zinc plastic genes between mines and coasts and randomization tests. size set 1 = Welsh population number of Differentially expressed genes control to salt/zinc; size set 2 = English population number of differentially expressed genes control to salt/zinc. Observed overlap = actual number of shared genes. Maximum randomised overlap = maximum number of shared genes from 10,000 randomisations. Empirical p value = frequency of observed overlap from 10,000 randomisations.

| Gene set of interest | size set 1 | size set 2 | observed overlap | maximum randomised overlap | empirical p value |
|---|---|---|---|---|---|
| Coastal shared salt plasticity | 2078 | 1676 | 957 | 191 | <0.0001 |
| Mine shared salt plasticity | 326 | 472 | 155 | 18 | <0.0001 |
| Coastal shared zinc plasticity | 13461 | 13343 | 10933 | 7918 | <0.0001 |
| Mine shared zinc plasticity | 241 | 836 | 143 | 23 | <0.0001 |

**Table S3.** Significantly enriched GO terms for shared Coastal salt-plastic genes only, with broader functional classification.

| ID | Description | broader function | gene number in GO | P value |
|---|---|---|---|---|
| GO:0010200 | response to chitin | stress response | 25 | 2.90E-09 |
| GO:0001101 | response to acid chemical | stress response | 90 | 8.40E-09 |
| GO:0009808 | lignin metabolic process | cell wall organisation | 16 | 2.10E-07 |
| GO:0120256 | olefinic compound catabolic process | metabolic processes | 10 | 7.20E-07 |
| GO:0006568 | tryptophan metabolic process | metabolic processes | 9 | 5.10E-06 |
| GO:0097305 | response to alcohol | stress response | 46 | 5.20E-05 |
| GO:0033993 | response to lipid | cell signalling | 63 | 0.00036 |
| GO:0042445 | hormone metabolic process | cell signalling | 25 | 0.00091 |
| GO:0060992 | response to fungicide | stress response | 4 | 0.00117 |
| GO:0009696 | salicylic acid metabolic process | stress response | 6 | 0.00144 |
| GO:0046351 | disaccharide biosynthetic process | metabolic processes | 6 | 0.00179 |
| GO:0009407 | toxin catabolic process | stress response | 6 | 0.0022 |
| GO:0046355 | mannan catabolic process | metabolic processes | 4 | 0.00254 |
| GO:0006032 | chitin catabolic process | stress response | 3 | 0.00415 |
| GO:0043312 | neutrophil degranulation | stress response | 3 | 0.00415 |
| GO:0019762 | glucosinolate catabolic process | metabolic processes | 4 | 0.00472 |
| GO:0032527 | protein exit from endoplasmic reticulum | other | 4 | 0.00472 |
| GO:0015718 | monocarboxylic acid transport | metabolic processes | 7 | 0.00614 |
| GO:0006879 | intracellular iron ion homeostasis | ion transport and homeostasis | 5 | 0.00615 |
| GO:0042759 | long-chain fatty acid biosynthetic process | metabolic processes | 6 | 0.00632 |
| GO:0006012 | galactose metabolic process | metabolic processes | 3 | 0.00639 |
| GO:0046686 | response to cadmium ion | stress response | 28 | 0.00663 |
| GO:0017001 | antibiotic catabolic process | stress response | 5 | 0.00688 |
| GO:0006820 | monoatomic anion transport | ion transport and homeostasis | 28 | 0.01125 |
| GO:0005978 | glycogen biosynthetic process | metabolic processes | 3 | 0.01267 |

| | | | | |
|---|---|---|---|---|
| GO:0006749 | glutathione metabolic process | stress response | 6 | 0.01434 |
| GO:0045332 | phospholipid translocation | other | 3 | 0.01675 |
| GO:2000379 | positive regulation of reactive oxygen species metabolic process | stress response | 3 | 0.01675 |
| GO:0009965 | leaf morphogenesis | growth and development | 11 | 0.01738 |
| GO:0055074 | calcium ion homeostasis | ion transport and homeostasis | 4 | 0.01786 |
| GO:0018105 | peptidyl-serine phosphorylation | other | 8 | 0.0202 |
| GO:0052545 | callose localization | cell wall organisation | 6 | 0.02025 |
| GO:0046890 | regulation of lipid biosynthetic process | metabolic processes | 4 | 0.02118 |
| GO:2000117 | negative regulation of cysteine-type endopeptidase activity | other | 3 | 0.02149 |
| GO:0000023 | maltose metabolic process | metabolic processes | 2 | 0.02419 |
| GO:0009805 | coumarin biosynthetic process | metabolic processes | 2 | 0.02419 |

**Table S4.** Significantly enriched GO terms for shared mine salt-plastic genes only, with broader functions classified.

| ID | Description | broader function | gene number in GO | P value |
|---|---|---|---|---|
| GO:0009414 | response to water deprivation | stress response | 10 | 0.00029 |
| GO:0009615 | response to virus | stress response | 5 | 0.00119 |
| GO:0048768 | root hair cell tip growth | growth and development | 2 | 0.00476 |
| GO:0009620 | response to fungus | stress response | 8 | 0.00883 |
| GO:0009664 | plant-type cell wall organization | cell wall organisation | 6 | 0.01598 |
| GO:0010192 | mucilage biosynthetic process | metabolic processes | 2 | 0.01714 |
| GO:0010119 | regulation of stomatal movement | stress response | 3 | 0.02015 |
| GO:0002376 | immune system process | stress response | 9 | 0.0242 |
| GO:0098869 | cellular oxidant detoxification | stress response | 4 | 0.03157 |
| GO:0010214 | seed coat development | growth and development | 2 | 0.03768 |
| GO:0009636 | response to toxic substance | stress response | 12 | 0.03823 |
| GO:0007568 | aging | other | 4 | 0.0411 |
| GO:0007187 | G protein-coupled receptor signaling pathway, coupled to cyclic nucleotide second messenger | cell signalling | 1 | 0.04321 |
| GO:0007188 | adenylate cyclase-modulating G protein-coupled receptor signaling pathway | cell signalling | 1 | 0.04321 |
| GO:0035461 | vitamin transmembrane transport | other | 1 | 0.04321 |
| GO:0071786 | endoplasmic reticulum tubular network organization | other | 1 | 0.04321 |
| GO:0098656 | monoatomic anion transmembrane transport | ion transport and homeostasis | 7 | 0.04453 |
| GO:0009860 | pollen tube growth | growth and development | 4 | 0.04734 |
| GO:0010150 | leaf senescence | other | 4 | 0.04734 |
| GO:0006468 | protein phosphorylation | other | 11 | 0.04992 |

**Table S5.** Significantly enriched GO terms shared across both coastal and mine ecotype, with broader functions classified.

| ID | Description | broader function | Coastal gene number in GO | Mine gene number in GO | Coastal P value | Mine P value |
|---|---|---|---|---|---|---|
| GO:0006970 | response to osmotic stress | stress response | 58 | 17 | 1.90E-07 | 4.50E-06 |
| GO:0009617 | response to bacterium | stress response | 52 | 12 | 6.20E-07 | 0.00278 |
| GO:0072722 | response to amitrole | stress response | 4 | 3 | 0.00022 | 2.20E-05 |
| GO:0010272 | response to silver ion | stress response | 5 | 3 | 0.00033 | 0.00018 |
| GO:0098542 | defense response to other organism | stress response | 61 | 18 | 0.00054 | 0.00153 |
| GO:0009635 | response to herbicide | stress response | 5 | 4 | 0.00072 | 7.00E-06 |
| GO:0016598 | protein arginylation | other | 3 | 1 | 0.00128 | 0.04321 |
| GO:0009828 | plant-type cell wall loosening | cell wall organisation | 4 | 2 | 0.00176 | 0.00399 |
| GO:0006979 | response to oxidative stress | stress response | 36 | 9 | 0.00183 | 0.0162 |
| GO:0009409 | response to cold | stress response | 32 | 7 | 0.00204 | 0.03899 |
| GO:0032412 | regulation of monoatomic ion transmembrane transporter activity | ion transport and homeostasis | 5 | 3 | 0.00688 | 0.00116 |
| GO:0010392 | galactoglucomannan metabolic process | metabolic processes | 3 | 2 | 0.01267 | 0.00328 |
| GO:0051070 | galactomannan biosynthetic process | metabolic processes | 3 | 2 | 0.01267 | 0.00328 |
| GO:0080167 | response to karrikin | cell signalling | 12 | 5 | 0.01913 | 0.00374 |

**Table S6.** Significantly enriched GO terms for cue transfer.

| ID | Description | number of genes in GO | P value |
|---|---|---|---|
| GO:0009635 | response to herbicide | 4 | 6.00E-10 |
| GO:0072722 | response to amitrole | 3 | 2.30E-08 |
| GO:0010272 | response to silver ion | 3 | 1.90E-07 |
| GO:0043312 | neutrophil degranulation | 2 | 1.80E-05 |
| GO:0006032 | chitin catabolic process | 2 | 1.80E-05 |
| GO:0009627 | systemic acquired resistance | 3 | 3.20E-05 |
| GO:0009615 | response to virus | 3 | 7.80E-05 |
| GO:0009620 | response to fungus | 4 | 0.00019 |
| GO:0007568 | aging | 3 | 0.00033 |
| GO:0010150 | leaf senescence | 3 | 0.00038 |
| GO:0051707 | response to other organism | 7 | 0.00111 |
| GO:0006955 | immune response | 5 | 0.00186 |
| GO:0006076 | (1->3)-beta-D-glucan catabolic process | 1 | 0.00592 |
| GO:0046902 | regulation of mitochondrial membrane permeability | 1 | 0.00691 |
| GO:0010262 | somatic embryogenesis | 1 | 0.00691 |
| GO:0015867 | ATP transport | 1 | 0.01083 |
| GO:0015866 | ADP transport | 1 | 0.01475 |
| GO:1901679 | nucleotide transmembrane transport | 1 | 0.01767 |
| GO:0009407 | toxin catabolic process | 1 | 0.02641 |
| GO:0006749 | glutathione metabolic process | 1 | 0.03794 |

**Table S7.** Significantly enriched GO terms for co-option.

| ID | Description | number of genes in GO | P value |
|---|---|---|---|
| GO:0006970 | response to osmotic stress | 9 | 3.10E-07 |
| GO:0009414 | response to water deprivation | 4 | 0.0025 |
| GO:0016598 | protein arginylation | 1 | 0.0099 |
| GO:0010243 | response to organonitrogen compound | 3 | 0.0121 |
| GO:0010726 | positive regulation of hydrogen peroxide metabolic process | 1 | 0.0138 |
| GO:0046244 | salicylic acid catabolic process | 1 | 0.0157 |
| GO:0060992 | response to fungicide | 1 | 0.0196 |
| GO:0015700 | arsenite transport | 1 | 0.0196 |
| GO:1902418 | (+)-abscisic acid D-glucopyranosyl ester transmembrane transport | 1 | 0.0216 |
| GO:0048316 | seed development | 4 | 0.0218 |
| GO:0046355 | mannan catabolic process | 1 | 0.0235 |
| GO:0010205 | photoinhibition | 1 | 0.0235 |
| GO:0010941 | regulation of cell death | 2 | 0.0237 |
| GO:0046685 | response to arsenic-containing substance | 1 | 0.0312 |
| GO:0009409 | response to cold | 3 | 0.0325 |
| GO:1901684 | arsenate ion transmembrane transport | 1 | 0.037 |
| GO:0042447 | hormone catabolic process | 1 | 0.0408 |
| GO:0017001 | antibiotic catabolic process | 1 | 0.0465 |

**Table S8.** Sample names and read counts (F+R) for salt experiment plants. RNA suffix indicates an individual plant sampled from living *S. uniflora* collections. Those with no suffix were more recently germinated for the experiment.

| Sample name | Plant name | raw read counts F+R | trimmed read counts F +R |
|---|---|---|---|
| SA02_C_7 | SA 2 | 42335250 | 41749450 |
| SA02_S_8 | SA 2 | 40844158 | 40318092 |
| SA04_C_15 | SA 4 | 41093634 | 40623636 |
| SA04_S_16 | SA 4 | 41137778 | 40681912 |
| SA07_C_11 | SA 7 | 40270096 | 39773232 |
| SA07_S_12 | SA 7 | 41422260 | 40905500 |
| GR-RNA-6_C_5 | GR RNA 6 | 43722884 | 43224348 |
| GR-RNA-6_S_6 | GR RNA 6 | 42254266 | 41660636 |
| GR-RNA-10_C_13 | GR RNA 10 | 42256298 | 41771490 |
| GR-RNA-10_S_14 | GR RNA 10 | 41263170 | 40859672 |
| GR-RNA-12_C_23 | GR RNA 12 | 42829958 | 42299184 |
| GR-RNA-12_S_24 | GR RNA 12 | 44029256 | 43489784 |
| BD03_C_17 | BD 3 | 43535140 | 43087694 |
| BD03_S_18 | BD 3 | 41198658 | 40741972 |
| BD05_C_9 | BD 5 | 41026254 | 40539942 |
| BD05_S_10 | BD 5 | 42056414 | 41528166 |
| BD07_C_19 | BD 7 | 41615692 | 41156374 |
| BD07_S_20 | BD 7 | 43001934 | 42450886 |
| PP1_C_3 | PP 1 | 41515228 | 40961716 |
| PP1_S_4 | PP 1 | 39593352 | 39066246 |
| PP12_C_21 | PP 12 | 40989006 | 40379610 |
| PP12_S_22 | PP 12 | 41973704 | 41435506 |
| PP-RNA-1_C_1 | PP RNA 1 | 41920488 | 41382040 |
| PP-RNA-1_S_2 | PP RNA 1 | 40856702 | 40400804 |

**Dataset S1 (SI_data_S1_ASP_annots.csv)**. Functional annotations for genes present in the shared coastal (ancestral) salt plasticity responses. Includes EggNOG orthologues, COG category, GO functions, Protein family names, and Kegg pathways. Produced using the *Silene uniflora* reference genome by Osborne *et al.* (10).

**Dataset S2 (SI_data_S2_DSP_annots.csv).** Functional annotations for genes present in the shared Mine (descendent) salt plasticity responses. Includes EggNOG orthologues, COG category, GO functions, Protein family names, and Kegg pathways. Produced using the *Silene uniflora* reference genome by Osborne *et al.* (10).

**Dataset S3 (SI_data_S3_cue_transfer_annots.csv).** Functional annotations for genes present in the cue transfer gene set. Includes EggNOG orthologues, COG category, GO functions, Protein family names, and Kegg pathways. Produced using the *Silene uniflora* reference genome by Osborne *et al.* (10).

**Dataset S4 (SI_data_S4_cooption_annots.csv).** Functional annotations for genes present in the co-option gene set. Includes EggNOG orthologues, COG category, GO functions, Protein family names, and Kegg pathways. Produced using the *Silene uniflora* reference genome by Osborne *et al.* (10).

**Dataset S5 (SI_data_S5_salt_gene_count_matrix.csv).** Raw gene count matrix generated in StringTie v2.2.0 (12) for salt experiment dataset.

**Dataset S6 (SI_data_S6_zinc_gene_count_matrix.csv).** Raw gene count matrix generated in StringTie v2.2.0 (12) for zinc experiment dataset.

**Dataset S7 (SI_data_S7_phenotype_data.xlsx).** Phenotype data from salt and zinc experiment samples used for input into DEseq2 datasets in R. Tab 1 is the salt experiment phenotype data, Tab 2 is the zinc experiment phenotype data and Tab 3 is the combined dataset for both experiments. The column Pop_treat shows the Population*treatment factor levels assigned for the first DEseq2 model structure. For combined experiment data, the salt control treatment was assigned the code C1 and zinc control was assigned the code C2.
